# EGFR Signalling Coordinates Patterning with Cell Survival During *Drosophila* Epidermal Development

**DOI:** 10.1101/399865

**Authors:** Samuel H. Crossman, Sebastian J. Streichan, Jean-Paul Vincent

## Abstract

Extensive apoptosis is often seen in patterning mutants, suggesting that tissues can detect and eliminate potentially harmful mis-specified cells. Here we show that the pattern of apoptosis in the embryonic epidermis of *Drosophila* is not a response to fate mis-specification but can instead be explained by the limiting availability of pro-survival signalling molecules released from locations determined by patterning information. In wild type embryos, the segmentation cascade elicits the segmental production of several EGFR ligands, including the TGF-alpha, Spitz and the Neuregulin, Vein. This leads to an undulating pattern of signalling activity, which prevents expression of the pro-apoptotic gene *hid* throughout the epidermis. In segmentation mutants, where specific peaks of EGFR ligands fail to form, gaps in signalling activity appear, leading to coincident *hid* upregulation and subsequent cell death. These data provide a mechanistic understanding of how cell survival, and thus appropriate tissue size, is made contingent on correct patterning.

## Introduction

Defective cells are often eliminated by apoptosis during development and tissue homeostasis (1-4). This has been particularly well studied during the process of cell competition, whereby unfit cells are eliminated when confronted with normal cells within a growing tissue (5). Excess apoptosis is also seen in mutants that lack essential developmental determinants, a phenomenon that has been observed in a variety of model organisms, including zebrafish embryos lacking the signalling molecule Sonic Hedgehog (6), mice lacking the negative Wnt signalling regulator APC in the developing neural crest (7) and *Drosophila* segmentation mutants (8-12). These observations have suggested the existence of a quality control system that detects conflicting or non-sense patterning inputs and, as a result, initiates apoptosis in response.

Although apoptosis was first observed in *Drosophila* segmentation mutants over thirty years ago (8-10), relatively little is known about the molecular basis of cell elimination in this context. Previous studies have revealed that apoptosis occurs within and around the areas where the missing developmental regulators are normally required. Thus, in the pair-rule mutant *odd-skipped* (*odd*), apoptosis is seen in alternate stripes that encompass the areas where *odd* is normally expressed (13, 14). Likewise, in embryos lacking the anterior determinant Bicoid, ectopic cell death occurs primarily at the anterior of the embryonic epidermis (13). In *Drosophila*, apoptosis is initiated by a double inhibition mechanism: the activity of ubiquitously expressed Inhibitor of Apoptosis Proteins (IAPs), which prevent caspase activation, is inhibited by the so-called IAP-antagonists *reaper, hid, grim* and *sickle* (15). Most apoptotic events are initiated by the transcriptional upregulation of one or more IAP-antagonist. Indeed, *hid* is upregulated in a pattern that prefigures apoptosis in segmentation mutants, i.e. at the anterior of *bicoid* mutants and in alternate stripes in *odd* mutants (13). It appears therefore that cells within the segmental pattern ‘know’ that they are missing key fate determinants and activate *hid* expression in response. Here we take advantage of the genetic tools in *Drosophila* to investigate how apoptosis is triggered in mis-patterned epidermal cells.

## Results

### Hid mediates apoptosis in *Drosophila* segmentation mutants

Consistent with an earlier report (13), extensive apoptosis was detected in Drosophila embryos lacking genes acting at various steps of the segmentation cascade, including mutants of the terminal gene *tailless* (*tll*), the gap gene *krüppel* (*kr*), the pair-rule gene *fushi-tarazu* (*ftz*), or the segment polarity gene *hedgehog* (*hh*) (Figure 1A-E). In each instance, cleaved Death Caspase-1 (Dcp1), a marker of apoptosis, was strongly enriched in the areas where the mutated patterning gene is known to be required during normal development. For example, in a null *ftz* mutant generated for this study (*ftz*^*Δ.attP*^), seven bands of apoptotic cells appeared in the segments where Ftz is normally expressed in the wild type (Figure 1E, see also (12)). Thus, in *ftz*^*Δ.attP*^ mutants, alternate segments undergo massive apoptosis, while almost no cell death is detected in the remaining, normally patterned, segments, which therefore serve as a useful control. For this reason, *ftz* mutants were selected as a prototypical condition to investigate mis-patterning-induced apoptosis.

**Fig. 1.**
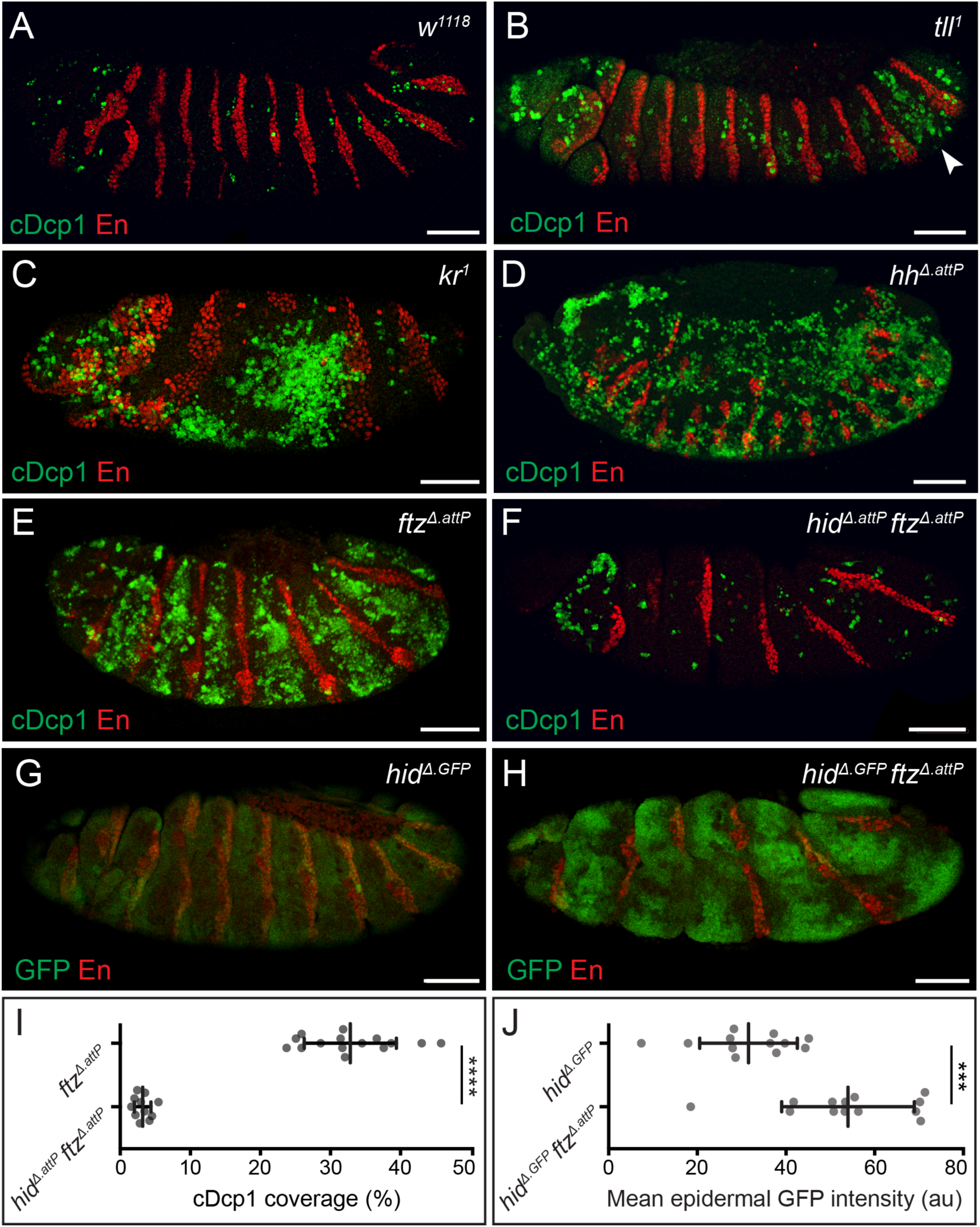
Widespread apoptosis in segmentation mutants is mediated by *hid*. (A-E) Cleaved Dcp1 immunoreactivity (cDcp1, green), in control (A), *tailless* (B), *krüppel* (C) *hedgehog* (D) and *fushi-tarazu* (E) embryos. Segmental enrichment of cDcp1 is detected around the regions where the mutated gene is known to act during normal development. Arrowhead in B indicates the zone of elevated Dcp1 cleavage in the posterior epidermis of *tll*^*1*^ embryos. Anti-Engrailed (En, red) provides a positional reference along the anteroposterior axis throughout. (F) Epidermal cDcp1 immunoreactivity in *hid*^*Δ.attP*^ *ftz*^*Δ.attP*^ double homozygotes is strongly reduced. (G, H) Transcription of *hid*, as assayed with the *hid*^*Δ.GFP*^ reporter, is detected at uniform low levels in a wild type background (G) but is upregulated in a banded pattern in *hid*^*Δ.GFP*^ *ftz*^*Δ.attP*^ double homozygotes. Scale bars 50µm. (I) Quantification of cDcp1 levels in *ftz*^*Δ.attP*^ single mutant and *hid*^*Δ.attP*^ *ftz*^*Δ.attP*^ double mutant embryos. (J) Quantification of mean epidermal GFP intensity values throughout the epidermis of *hid*^*Δ.GFP*^ single mutant and *hid*^*Δ.GFP*^ *ftz*^*Δ.attP*^ double mutant embryos. In graphs, means are shown and error bars display standard deviation, *** = p < 0.001, unpaired Student’s t-test.

Among the four known pro-apoptotic genes of *Drosophila*, *hid* is the key mediator of apoptosis in *bicoid* mutant embryos (13). To assess the involvement of *hid* in *ftz* mutants, we created a clean *hid* allele (*hid*^*Δ.attP*^) by replacing the first coding exon, which encodes the IAP-binding motif, with an *attP* integrase site (16-18). This mutation was recombined with *ftz*^*Δ.attP*^ and the resulting double mutant embryos were stained with anti-cleaved Dcp1. Although some signal remained in the head region, immunoreactivity was much lower than in single *ftz*^*Δ.attP*^ mutants (Figure 1F, I). In contrast, abundant apoptosis was detected in *ftz*^*Δ.attP*^ mutants lacking the closely related IAP-antagonists, *reaper* and *sickle* (Figure S1). We conclude that, as in *bicoid* mutants, *hid* is the main mediator of apoptosis in *ftz* mutant embryos.

To visualize the expression of *hid* in mis-patterned embryos, we generated an authentic transcriptional reporter by integrating a cDNA encoding eGFP into the attP site of *hid*^*Δ.attP*^ (16). These animals will be referred to as *hid*^*Δ.GFP*^ to indicate that this genetic modification creates a null allele (Δ) as well as a fluorescent reporter (GFP). In homozygous *hid*^*Δ.GFP*^ embryos, weak GFP signal could be detected throughout the epidermis, suggesting that low level *hid* expression occurs at sub-apoptotic levels during normal development (Figure 1G). In *hid*^*Δ.GFP*^ *ftz*^*Δ.attP*^ double homozygotes, increased levels of GFP fluorescence were detected in broad stripes resembling the bands of caspase immunoreactivity seen in *ftz*^*Δ.attP*^ single mutants (Figure 1H, J). Time-lapse imaging showed that striped GFP fluorescence arises around stage 11 of embryogenesis (Mov S1), approximately 4 hours after the time when pair-rule genes contribute to segmental patterning (19, 20). Likewise, cleaved Dcp1 immunoreactivity became only detectable at stage 11 in homozygous *ftz*^*Δ.attP*^ embryos (Figure S2A-C). Therefore, loss of Ftz appears to trigger *hid* expression and apoptosis with a delay, well after Ftz has fulfilled its patterning role in wild type embryos.

### EGFR signalling and *hid* expression are anti-correlated in the embryonic epidermis

Mismatched or non-sense cell fates could conceivably be recognized through local cell interactions. With the aim of identifying the relevant mediator, we used the ubiquitous *actin5C*-Gal4 driver to overexpress modulators of major signalling pathways in a *ftz*^*Δ.attP*^ mutant background and stained the resulting embryos with anti-cleaved Dcp1 (summarized in Table S1). One manipulation, overexpression of an activated form of the *Drosophila* EGF receptor (*EGFR*^*act*^, (21)), significantly reduced cleaved Dcp1 immunoreactivity (Figure 2A-D). Since EGFR signalling is required for survival of the cells that secrete naked cuticle (22) and since EGFR signalling has been previously implicated in suppressing *hid* activity (23, 24), we set out to determine whether the excess apoptosis and upregulation of *hid* in *ftz* mutants could be accounted for by a loss of EGFR activity.

**Fig. 2.**
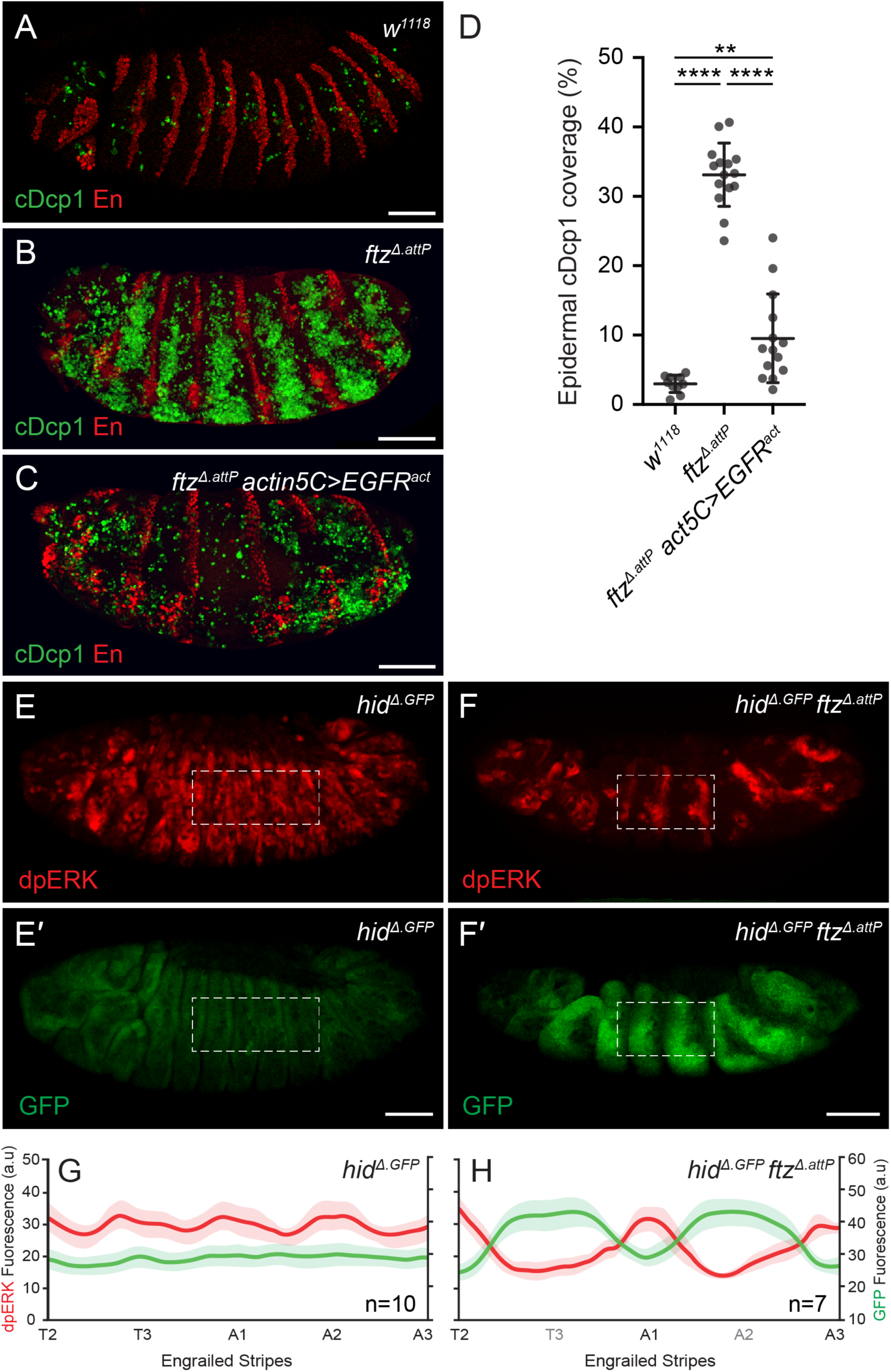
Loss of EGFR signalling and *hid* upregulation in *ftz* mutants (A-D) Cleaved Dcp1 immunoreactivity in control *w1118* (A, A′), *ftz*^*Δ.attP*^ (B, B′) and *ftz*^*Δ.attP*^ *actin5C*-Gal4, UAS-*EGFR* (C, C′) embryos. (D) Quantification of mean Dcp1 levels for genotypes shown in A-C. Error bars display standard deviation, *** = p < 0.001, ** = p < 0.01, unpaired Student’s t-test. (E, F) dpERK/GFP immunoreactivity in *hid*^*Δ.GFP*^ single mutant (E, E′) and *hid*^*Δ.GFP*^ *ftz*^*Δ.attP*^ double mutant embryos (F, F′). Dashed line indicates representative region of interest for quantification in panels G and H. Scale bars 50µm. (G, H) Mean dpERK (red) and GFP (green) fluorescence intensity profiles in control *hid*^*Δ.GFP*^ (G) and *hid*^*Δ.GFP*^ *ftz*^*Δ.attP*^ (H) samples. Engrailed expression stripes were used as a spatial marker along the A/P axis (see material and methods). Shaded areas depict S.E.M.

If EGFR signalling is needed to keep *hid* expression off, we would expect *hid* expression to be upregulated in EGFR mutants. Indeed, strong ubiquitous GFP fluorescence was observed in embryos homozygous for both *EGFR*^*F3*^, an amorphic allele, and *hid*^*Δ.GFP*^ (Figure S3A, B). This suggests that most, if not all, epidermal cells require EGFR signalling to keep *hid* expression low. Conversely, the pro-survival activity of EGFR signalling appears to be largely mediated by repression of *hid* activity since almost no apoptosis was detected in *EGFR*^*F3*^ *hid*^*Δ.attP*^ double mutants (Figure S3D). In *EGFR*^*F3*^ single mutant embryos, cell death was widespread (Figure S3C) but did not occur until embryonic stage 11 (Figure S2D-F), the same time when apoptosis becomes detectable in *ftz* mutants. We conclude that, in the absence of EGFR signalling, most cells of the epidermis upregulate *hid* expression and undergo apoptosis around mid-embryogenesis.

To visualize EGFR signalling activity in mis-patterned *ftz* embryos, we stained embryos with an antibody that recognizes phosphorylated extracellular signal-regulated kinase (dpERK, (25)). As validation, we first stained *EGFR*^*F3*^ homozygotes. No immunoreactivity was detectable above background levels (Figure S4), a strong indication that EGFR is the principal contributor of ERK phosphorylation in the embryonic epidermis. Anti-dpERK was next used to compare the patterns of EGFR signalling activity in *ftz*^*Δ.attP*^ mutant and control embryos. The *hid*^*Δ.GFP*^ allele (homozygous) was included to allow simultaneous assessment of *hid* expression and to avoid possible confounding effects of apoptosis on signalling activity. In control *hid*^*Δ.GFP*^ embryos, dpERK immunoreactivity was detectable in nearly all epidermal cells, though not uniformly (Figure 2E). Signal intensity followed an undulating pattern along the anterior-posterior (A/P) axis, with smooth peaks near segment boundaries and intervening shallow troughs (Figure 2G). Little GFP fluorescence was detectable in the epidermis (Figure 2E′, G), confirming that *hid* transcription (as reported by *hid*^*Δ.GFP*^) remains low in normally patterned embryos (see also Figure 1G & S3A). In the absence of *ftz* activity, dpERK immunoreactivity dropped significantly in seven broad bands (Figure 2F, H), leaving only half the number of signalling peaks intact. At the same time, a complementary pattern of GFP fluorescence (from the *hid*^*Δ.GFP*^ reporter) appeared (Figure 2F, H). We conclude that, in *ftz* mutant embryos, the transcriptional activation of *hid* increases in regions where EGFR signalling activity fails to reach the threshold level required for epidermal cell survival.

Other segmentation mutants, including *tll*, *kr*, and *hh* also displayed anti-correlated *hid*^*Δ.GFP*^ reporter activity and dpERK immunoreactivity (Figure 3A-C). Importantly, the pattern of *hid* expression in these mutants corresponded to that of apoptosis (as shown in Figure 1A-D). Embryos lacking another segmentation gene, *patched*, were particularly informative as they had increased dpERK immunoreactivity and did not upregulate *hid*^*Δ.GFP*^ (Figure 3D). Crucially, they also showed no excess apoptosis, as assayed with anti-cleaved Dcp1 (Figure S5). This observation confirms the tight correlation between EGFR signalling and the absence of *hid* expression. It also shows that not all instances of patterning error are associated with apoptotic cell death.

**Fig. 3.**
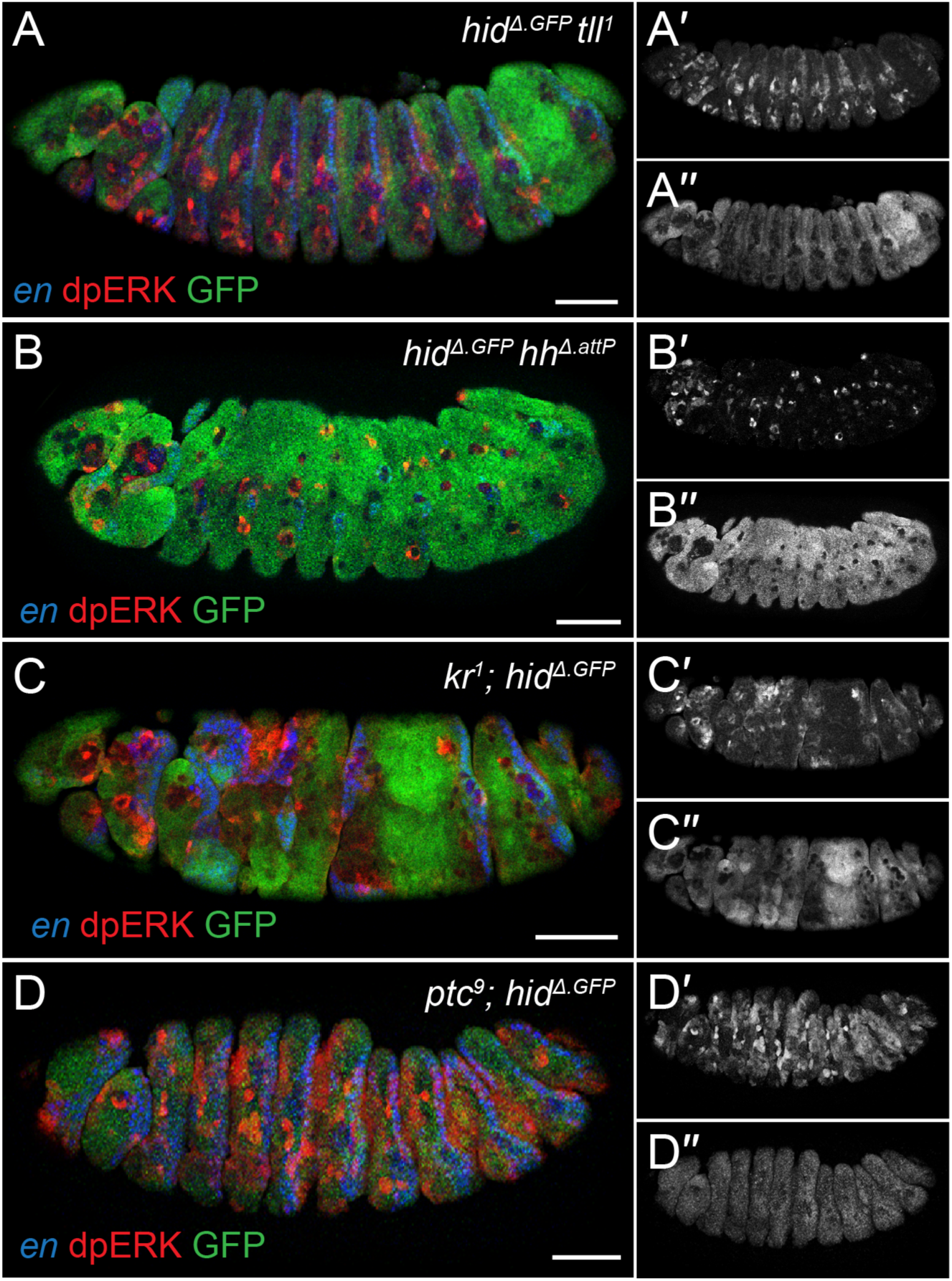
*hid* and dpERK are anti-correlated in various patterning mutants. (A-D) GFP and dpERK immunoreactivity in *hid*^*Δ.GFP*^ *tll*^*1*^ (A), *hid*^*Δ*.*GFP*^ *hh*^*Δ.attP*^ (B), *kr*^*1*^*; hid*^*Δ.GFP*^ and *ptc*^*9*^*; hid*^*Δ.GFP*^ (D) double mutants. In each instance, dpERK intensity and *hid*^*Δ.GFP*^ expression are anti-correlated. Scale bars 50µm.

### Segmentally expressed EGF ligands coordinate apoptosis with the patterning cascade

EGFR signalling has previously been widely implicated in cell survival in the embryo and other tissues (22, 26, 27). Known *Drosophila* EGFR ligands include the TGF-α homologues Spitz, Keren and Gurken, which require activation by Rhomboid proteases (28-31), as well as the Rhomboid-independent neuregulin-like ligand Vein (32). To assess the nature of the ligands involved in the survival of embryonic epidermal cells, we first examined embryos homozygous for a deficiency (Df ^*rho*-1,3^) spanning the *rhomboid* (*rho*-1) and *roughoid* (*rho*-3) loci. In this background, a relatively mild apoptotic phenotype was observed (Figure S6B). Likewise, homozygous *vein* mutants were also characterized by a relatively minor increase in the number of apoptotic cells compared to wild type embryos (Figure S6C). In contrast, simultaneous removal of both ligand systems, in Df ^*rho*-1,3^ *vn*^*L6*^ double homozygotes, led to widespread apoptosis (Figure S6D), consistent with the severe cuticle phenotype previously reported (22). We conclude that multiple ligands act redundantly to elicit the EGFR signalling landscape that, at mid-embryogenesis, prevents apoptosis.

In agreement with the observed pattern of EGFR signalling, *vein* and *rho-1*, were found to be expressed in a generally segmental manner in wild type embryos (Figure 4A, C). Thus, peaks of ligand production correspond roughly to the areas of high dpERK immunoreactivity seen around segmental borders (Figure 2G). We therefore considered whether changes in ligand production could account for the observed changes in EGFR signalling in patterning mutants. The pattern of *rho-1* expression is expected to depend on the segmentation network since it is determined by segment polarity genes (33, 34), which are themselves regulated by upstream pair-rule and gap genes. It is likely that the segmentation network similarly controls *vein* expression. As expected therefore, peaks of *vein* and *rho-1* expression failed to appear in alternate segments of *ftz*^*Δ.attP*^ mutants (Figure 4B, D), foreshadowing the loss of dpERK immunoreactivity (Figure 2F), *hid* upregulation (Figure 1H), and apoptosis (Figure 1E). Importantly, the same logic can explain the pattern of apoptosis in other segmentation mutants, which are all expected to affect the pattern of EGFR ligand production within their realm of action. It also neatly explains the absence of apoptosis in *ptc* mutants, where ectopic peaks of *rho-1* expression arise (33).

**Fig. 4.**
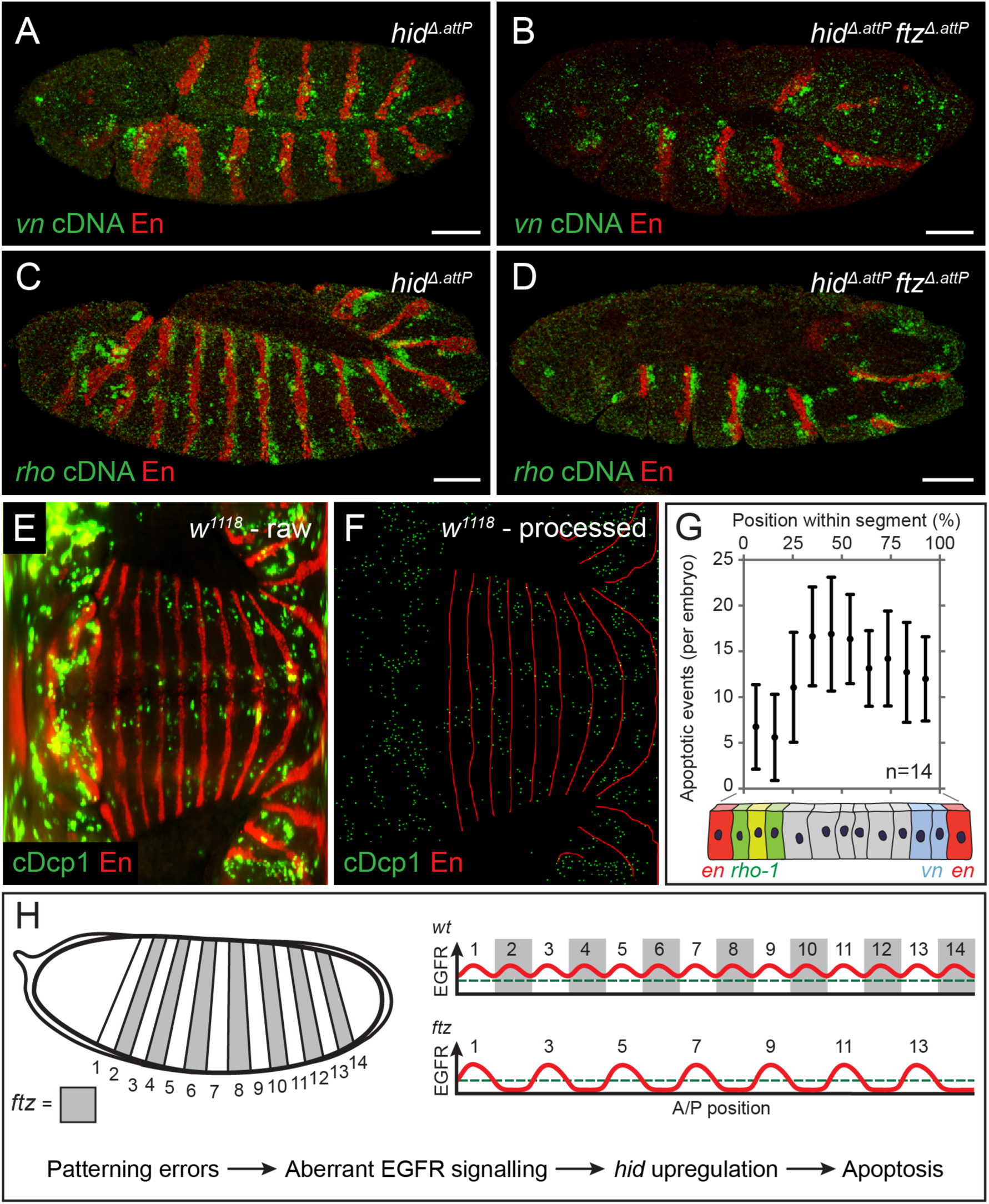
Patterned expression of *vein* and *rhomboid* ensure cell survival (A, B) Expression of *vn* in control *hid*^*Δ.attP*^ (A) and *hid*^*Δ.attP*^ *ftz*^*Δ.attP*^ (B) homozygotes, as detected by fluorescent *in situ* hybridisation. *hid*^*Δ.attP*^ is included in the background to avoid any confounding effects of apoptosis (C, D) Expression of *rho-1* in *hid*^*Δ.attP*^ (C) and *hid*^*Δ.attP*^ *ftz*^*Δ.attP*^ homozygotes, as detected by fluorescent *in situ* hybridisation. Scale bars 50µm. (E) Representative raw mSPIM image of a *w*^*1118*^ embryo (control) stained with anti-cDcp1 and anti-Engrailed (En, red). Anterior is to the left and posterior to the right, with the ventral midline running horizontally in the middle. (F) Processed version of the image shown in A. cDcp1-positive cells were segmented and reduced to individual pixels (green). Similarly, stripes of *engrailed* expression were reduced to a skeleton (red) to allow the position of each apoptotic cell to be mapped along the A/P axis of the segment and thus to derive a relative position applicable to all segments. (G) Histogram displaying the frequency of apoptosis along the A/P axis of the segment. Illustration of the en, rho-1 and vn expression domains is included to indicate the sites of ligand production. Error bars show standard deviation. (H) Model explaining the pattern of apoptosis in *ftz* and other segmentation mutants. In normal development, the segmental expression of EGFR ligands maintains signalling activity above the threshold level required for cell survival (green dashed line). When patterning errors disrupt the landscape of EGFR activity, signalling falls below the survival threshold, triggering *hid* upregulation and patterned apoptosis.

During normal development, EGFR signalling is lowest in cells located furthest away from a source of ligand (Figure 2G). We asked whether these cells are particularly susceptible to apoptosis. To map the pattern of apoptosis along the A/P axis, we performed *in toto* multiview single plane illumination microscopy (mSPIM) on wild-type embryos stained with anti-cleaved Dcp1 and anti-Engrailed, which marks the anterior edge of each segment boundary. This approach allowed us to visualize, at a given time point, all epidermal apoptotic events, which are relatively rare during normal development (see Figure 1A and 2A). Raw image files were generated (Figure 4E), segmented (Figure 4F) and the position of each caspase positive cell was then mapped and normalized relative to the nearest anterior and posterior segment boundary. These data were then tabulated in a histogram displaying the frequency of apoptosis at various positions along the anterior-posterior axis within each segment (Figure 4G). Cell death was relatively infrequent near segment boundaries, where *vein* and *rhomboid* are produced, while apoptosis was more abundant in the middle of the intervening areas. We conclude that, in the cells that are most distant from the source of ligand, EGFR signalling activity falls near, and occasionally below, the threshold level needed to protect against apoptosis. Cells in these regions are therefore more likely to commit to an apoptotic cell fate.

## Discussion

Excess apoptosis in patterning mutants gives the impression that mis-patterned cells are recognized as deleterious and eliminated during development. We have shown that apoptosis in embryonic patterning mutants can be explained by the requirement of EGFR signalling for cell survival and the fact that the segmentation cascade culminates in the segmental production of EGF ligands (model summarized in figure 4D). Thus, in segmentation mutants, sources of EGF ligand production are lost, leaving remaining sources unable to reach all intervening cells. This results in tissue loss until all cells are brought back within signalling range, restoring a ‘normal’ distance between segment boundaries. EGFR signalling has previously been linked to compartment size regulation in the embryo (35). Specifically, it was proposed that epidermal cells in the posterior regions of each segment compete for a finite amount of Spitz emanating from an anterior source. Our results are consistent with an extended version of this model, whereby multiple EGFR ligands (including Spitz and Vein) produced at both sides of each segment boundaries control the size of whole segments, not just posterior compartments. The pattern of apoptosis in wild type embryos (Figure 4G) suggests that each segment is slightly oversized upon completion of patterning and that the extra cells are eliminated as a result of sub-threshold EGFR signalling and *hid* activation. Accordingly, there is no need to invoke the existence of a mechanism that recognizes and eliminates mis-specified cells, and mis-patterning-induced cell death could be the byproduct of a size control system. It is worth noting that the lack of EGFR signalling triggers *hid* expression only at stage 11, when proliferation is largely complete (36). This would serve as a particularly suitable time for a size control checkpoint and pruning of excess cells would be an effective means of ensuring reproducible segment dimensions in a fast-growing embryo with no compensatory proliferation. Similar pro-survival signalling by limited ligand diffusion from specific locations could form the basis of tissue size sensing, and explain mis-patterning-induced apoptosis, in a variety of developmental contexts.

## Materials and methods

### *Drosophila* stocks and husbandry

Flies were maintained at 25°C on standard fly food containing yeast, agar and cornmeal. *w*^*1118*^ was used as a wild type control throughout.

To identify regulators of apoptosis in mis-patterned cells, a collection of UAS-lines was assembled. Each expressed a gene that modulates the activity of a signalling pathway in the presence of Gal4 (Table S1). These so-called UAS lines were crossed to the null *ftz* allele *ftz*^*Δ.attP*^ and male progeny containing both the ftz allele and the UAS transgene were crossed to females bearing the ubiquitous *actin5C*-Gal4 driver recombined to *ftz*^*Δ.attP*^. The embryos produced from this cross were then stained for cleaved-Dcp1.

### Generation of transgenic flies

Accelerated ‘ends out’ homologous recombination was used to generate the *hid*^*Δ.attP*^ and *ftz*^*Δ.attP*^ alleles (as described in (18)). Homology arms of approximately 2kb were PCR amplified from genomic DNA. These fragments were cloned into a targeting vector (pTV^Cherry^) containing a GMR>*white* eye marker and an attP landing site flanked with FRT sites. UAS-*reaper* was included downstream of the 3’ homology arms to favour successful homologous recombination. Targeting vectors were integrated into the genome via P-element mediated transformation to generate so-called donor lines. To release the homology cassette from it genomic location, donor flies were crossed to *hs*-FLP and heat shocked at 24hr intervals throughout larval development. Adult female progeny with mottled eye colour (indicating the presence of both the targeting vector and hs-FLP constructs) were crossed to the strong *ubi*-Gal4 driver to eliminate individuals that had either failed to excise the homology cassette or had undergone an unsuccessful recombination event. Resultant *white*^*+*^ candidates were then isolated and verified with PCR.

To generate the *hid*^*Δ.GFP*^ reporter, an eGFP cDNA was cloned into the RIV^Cherry^ vector (18), which contains an attB sequence and a *3xPax3-cherry* selection marker. This construct was injected into *hid*^*Δ.attP*^ embryos along with a source of ΦC31 integrase and progeny were screened for Cherry expression. Rainbow Transgenic Flies Inc (Camarillo, CA, USA) and Bestgene Inc (Chino Hills, CA, USA) were used for embryo injection services.

### Embryo collection

Overnight embryo collections were transferred to baskets and washed with water. Embryos were then dechorionated using 75% bleach for 3 minutes and washed again. After drying on a paper towel, embryos were transferred with a brush to microcentrifuge tubes containing equal volumes of heptane and 6% formaldehyde in phosphate buffered saline (PBS). Fixation was performed for 25 minutes on a rotating platform. The fixative was then removed and replaced with 1ml methanol. Embryos were shaken vigorously for 45 seconds to remove the vitelline membrane and the heptane was removed. Embryos were washed twice more in methanol and moved to - 20°C for long-term storage.

### Immunofluorescence

Fixed embryos were rehydrated in 0.3% triton in PBS (PBT) and blocked for 30 minutes at room temperature in 4% bovine serum albumin in PBT. Embryos were incubated with primary antibody at 4°C overnight, washed for 1 hour in PBT at room temperature and then incubated in a solution of Alexa Fluor 488, 568, or 633 conjugated species appropriate secondary antibodies (Thermo Fisher Scientific, 1:1000) for 2 hours. Further washes (1hour) were conducted before samples were mounted in Vectashield containing DAPI (Vector laboratories). Primary antibodies used in this study are listed below.

### Confocal microscopy

Confocal microscopy was conducted with a Leica SP5 confocal microscope using a 20x (Leica, NA 0.7) or 40x (Leica, NA 1.25) oil immersion objective. Typically, 8 confocal planes were imaged at 1.8µm intervals and processed using Fiji (NIH) and Photoshop (Adobe).

### Live confocal imaging

Samples were prepared using the hanging drop method described in (37). A small drop of oxygen permeable Voltalef 10S oil was spotted onto a cover slip. A dechorionated embryo was then placed inside the oil and oriented with a 27G hypodermic needle. The sample was then inverted so that the oil containing the sample hung below the cover slip. Images were acquired from above at 10 minute intervals using a Leica SP5 confocal microscope and processed using imageJ (NIH) and Photoshop (Adobe) as described above.

### mSPIM imaging

Fluorescently labelled samples where imaged on a multi view light sheet microscope (38). The optics consisted of two detection and illumination arms. Here each detection arm formed an epifluorescence microscope using in sequence: A water-dipping lens (Apo LWD 25x, NA 1.1, Nikon Instruments Inc.), a filter wheel (HS-1032, Finger Lakes Instrumentation LLC) equipped with emission filters (BLP01-488R-25, BLP02-561R-25, Semrock, Inc), a tube lens (200 mm, Nikon Instruments Inc.), and sCMOS camera (Hamamatsu Flash 4.0 v2.0). The imaging produced an effective pixel size of 0.26 µm. The illumination arms each consisted in sequence: a water-dipping objective (CPI Plan Fluor 10x, NA 0.3, Nikon Instruments Inc), a tube lens (200 mm, Nikon Instruments Inc) a scan lens (S4LFT0061/065, Sill optics GmbH & Co. KG), and a galvanometric scanner (6215h, Cambridge Technology Inc.). The illumination arms where fed by a combination of lasers (06-MLD 488nm, Cobolt AB, and 561 LS OBIS 561nm, Coherent Inc.). Samples where translated through the resulting lightsheet, using a linear piezo stage (P-629.1CD together with E-753 controller). Multiple rotation views where achieved using a rotational piezo stage (U-628.03 with C-867 controller), in combination with a linear actuator (M-231.17 with C-863 controller, all Physik Instrumente GmbH and Co. KG). Microscope operation was done using Micro Manager (39), running on a Super Micro 7047GR-TF Server, with 12 Core Intel Xeon 2.5 GHz, 64 GB PC3 RAM, and hardware Raid 0 with 7 2.0 TB SATA drives. 4 views with a rotational offset of 90^0^ where recorded at 1 µm sectioning. Fusion of individual views is based on a diagnostic specimen used to determine an initial guess for an affine transformation that maps each view into a common frame of reference. Together with the data, this initial guess was used with a rigid image registration algorithm (40) to fuse individual views and resulting in isotropic resolution of .26 µm in the final registered image. ImSAnE (41) was used to obtain tissue cartographic projections showing global maps of the embryo surface of interest.

### Fluorescent *in situ* hybridisation

Digoxigenin labelled antisense RNA probes were synthesized from Gold Collection (BDGP) clones containing full length versions of the *vein* or *rhomboid* cDNAs (clone number LP21849 and LD06131 respectively). Vectors were linearized with the EcoRI restriction enzyme and used as a template for *in vitro* transcription as per manufacturer instructions (Roche, SP6/T7 DIG RNA Labelling kit). Fixed embryos were bathed in 3% hydrogen peroxide in methanol for 15 minutes to remove endogenous peroxidase activity and rinsed extensively in PBT. Embryos were then incubated for 3 minutes in 10µg/ml proteinase K in PBT and washed in 2mg/ml Glycine in PBT. Samples were next washed in hybridization buffer before the relevant probe was added and the samples incubated overnight at 55°C. The next day, the embryos were washed once more in PBT and incubated overnight at 4°C with a sheep anti-DIG antibody diluted at 1:2000. On the third day, embryos were washed and incubated with a biotinylated anti-sheep secondary antibody and the final signal was developed using a Tyramide-FITC signal amplification kit (Thermo Fisher Scientific). Standard immunostaining was then performed on these samples, using the protocol outlined above.

### Quantification and statistical analysis

To measure fluorescence intensity along the A/P axis, rectangular regions of interest with a fixed height of 75µm were drawn over four contiguous segments. In each instance efforts were made to ensure the same embryonic regions were analyzed between samples. GFP (from *hid*^Δ.GFP^), dpERK and Engrailed intensity profiles were generated across the regions of interest in Fiji (42) and these data were imported into MATLAB (Mathworks) for further analysis. GFP and dpERK intensity profiles were fitted to the peaks of *engrailed* expression (i.e. segment boundaries) along the A/P axis to normalize data to segment width (which can vary between samples). This was achieved by creating 50 evenly spaced intervals along the A/P axis of each segment before extracting the raw data values and averaging the resulting numbers across all samples analyzed. SEM was calculated for each point and data was plotted using the boundedline.m function developed by Kelly Kearney and downloaded from the MATLAB stack exchange.

To measure the frequency of apoptosis along the segmental A/P axis, the Engrailed and cleaved-Dcp1 channels were separated from the raw mSPIM image files and then segmented using the open source iLastik software (University of Heidelberg). Segmented images were imported into ImageJ and reduced to single pixel lines and points using the software’s inbuilt skeletonisation algorithms. From these skeletons, individual segment boundaries were manually classified and masks were generated that covered each of the segments to be analyzed. Processed files were then transferred to MATLAB for further analysis. Distances were calculated from each Dcp-1 positive pixel to the nearest anterior and posterior Engrailed positive pixel using a k-nearest neighbours algorithm and these measurements were used to determine the position of each apoptotic cell along the segmental A/P axis as a percentage of segment width. Cells located in the immediate vicinity of the ventral midline (defined as 20% of linear Dorsal to Ventral distance) were excluded from the analysis to avoid apoptotic figures in the developing nervous system. The retained data were collated and tabulated as a histogram.

| *Drosophila* stocks used in the present study |  |  |
| --- | --- | --- |
| Genotype | Source | Identifier |
| *actin5C-GAL4/TM6B* | BDSC | FBst0003954 |
| *Df<sup>(3L)BSC178</sup>/TM6B (Df<sup>rho-1, 3</sup>)* | BDSC | FBst0009609 |
| *Df<sup>(3L)XR38</sup>/TM3* | Irene Miguel-Aliaga | N/A |
| *EGFR<sup>F3</sup>/CyO* | BDSC | FBst0034043 |
| *ftz<sup>Δ.attP</sup>/TM3* | This study | N/A |
| *hh<sup>Δ.attP</sup>/TM3* | Cyrille Alexandre (18) | N/A |
| *hid<sup>Δ.attP</sup>/TM3* | This study | N/A |
| *hid<sup>Δ.GFP</sup>/TM3* | This study | N/A |
| *kr<sup>1</sup>/CyO* | BDSC | FBst0003494 |
| *ptc<sup>9</sup>/CyO* | BDSC | FBst0003377 |
| *tll<sup>1</sup>/TM3, Sb* | BDSC | FBst0002729 |
| *UAS-tkv<sup>sct</sup>* | Markus Affolter | N/A |
| *UAS-hh* | Peter Lawrence | N/A |
| *UAS-EGFR<sup>act</sup>* | BDSC | FBst0009534 |
| *UAS-p53<sup>DN</sup>* | BDSC | FBti0040583 |
| *UAS-puc* | Enrique Martin-Blanco | N/A |
| *UAS-upd* | David Strutt (43) | FBtp0011049 |
| *UAS-wg* | Xavier Franch-Marro | N/A |
| *UAS-yki.GFP* | Nic Tapon | N/A |
| *vn<sup>L6</sup>/TM3* | Amanda Simcox | N/A |
| *w<sup>1118</sup>* | BDSC | FBst0003605 |

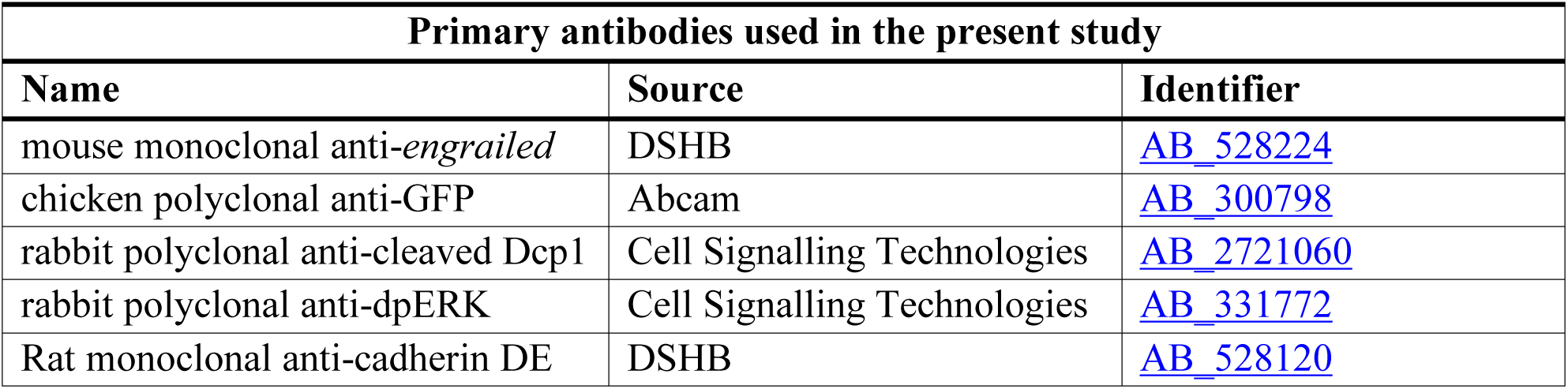

| Primary antibodies used in the present study |  |  |
| --- | --- | --- |
| Name | Source | Identifier |
| mouse monoclonal anti-*engrailed* | DSHB | AB_528224 |
| chicken polyclonal anti-GFP | Abcam | AB_300798 |
| rabbit polyclonal anti-cleaved Dcp1 | Cell Signalling Technologies | AB_2721060 |
| rabbit polyclonal anti-dpERK | Cell Signalling Technologies | AB_331772 |
| Rat monoclonal anti-cadherin DE | DSHB | AB_528120 |

## Acknowledgments

We thank Alberto Baena-Lopez for sharing fly stocks and reagents and Donald Bell for imaging support. We also acknowledge the advice and help of Ingrid Poernbacher, Kristina Stapornwongkul, Alberto Baena-Lopez and Jorge Beira during the early phase of this research. We also thank Barry Thompson, Ingrid Poernbacher and Ian McGough for comments on the manuscript. This work was supported by core funding (FC001204) from The Francis Crick Institute and an Investigator Award from the Wellcome Trust (206341/Z/17/Z).

## Author contributions

SHC and JPV designed the project and SHC performed the experiments. SJS performed mSPIM imaging and SHC analyzed and prepared the results. SHC and JPV wrote the manuscript.

## Figures and legends

**Fig. S1.**
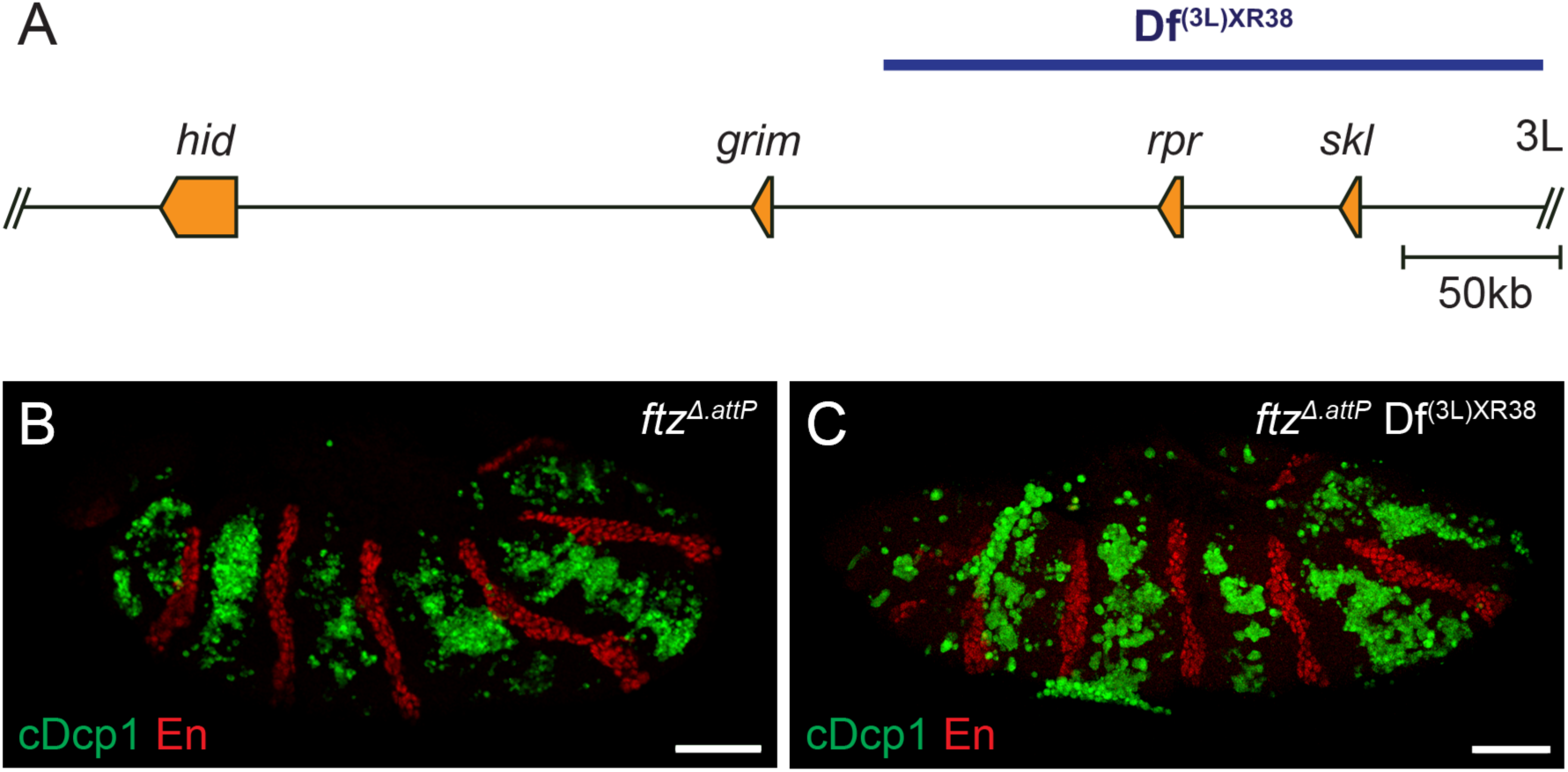
Apoptosis in *ftz* mutants does not require *reaper.* (A) Schematic representation of the *Drosophila* H99 locus showing the genes encoding the four major IAP antagonists. The Df^(3L)XR38^ deficiency, which removes the *rpr* and *skl* genes, is highlighted in blue. (B,C) Cleaved Dcp1 immunoreactivity in *ftz*^*Δ.attP*^ (B) and *ftz*^*Δ.attP*^ Df^(3L)XR38^ homozygotes (C). Scale bars 50µm.

**Fig. S2.**
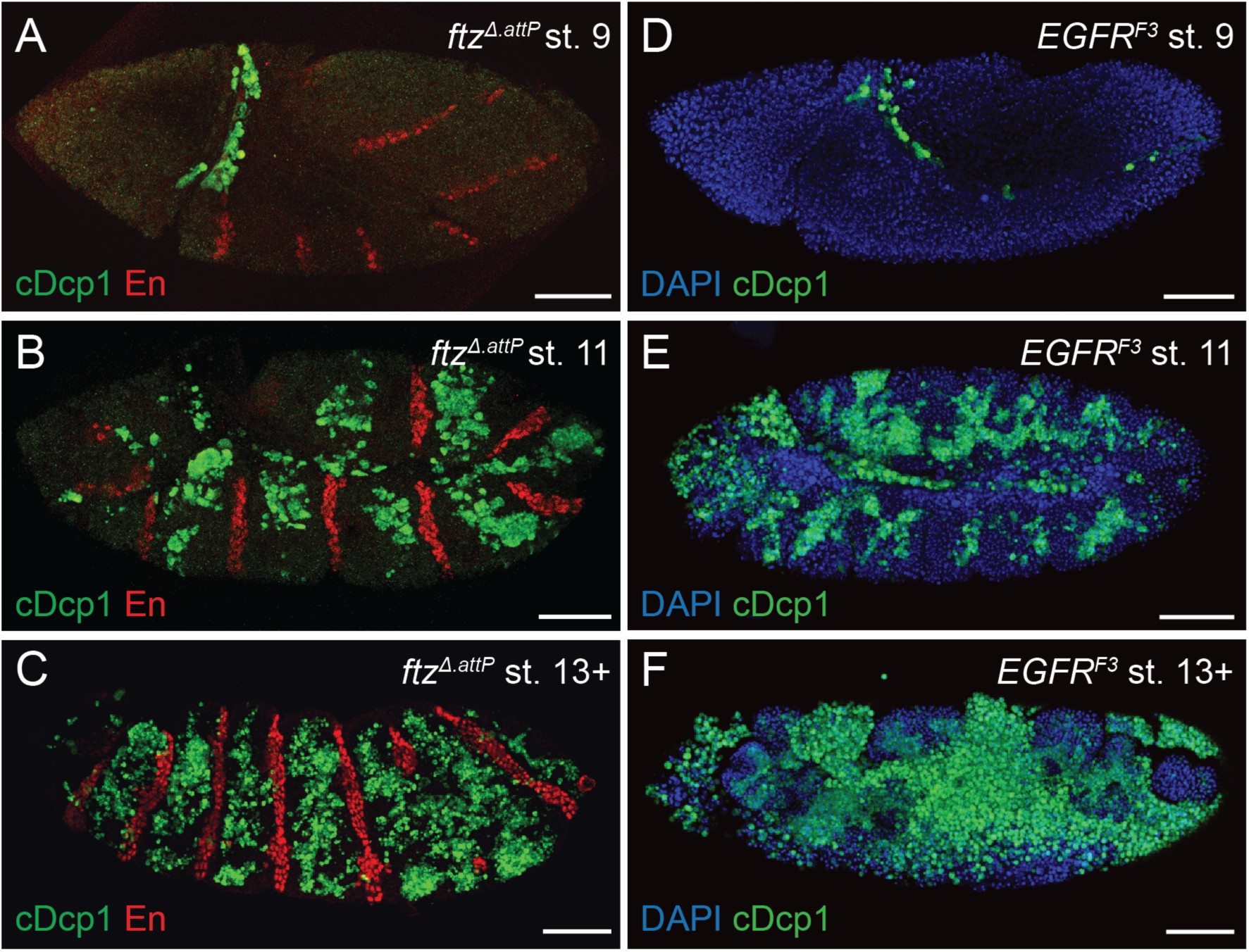
Apoptosis in *ftz* and *EGFR* mutants is first detectable at embryonic stage 11. (A-C) Cleaved Dcp1 immunoreactivity in *ftz*^*Δ.attP*^ mutants at embryonic stages 9 (A), 11 (B) and 13+ (C). (D-F) Cleaved Dcp1 immunoreactivity in *EGFR*^*F3*^ mutants at embryonic stages 9 (D), 11 (E) and 13+ (F). In both genotypes, cleaved Dcp1 is first detected in stage 11 and persists throughout the remainder of embryonic development. Scale bars 50µm.

**Fig. S3.**
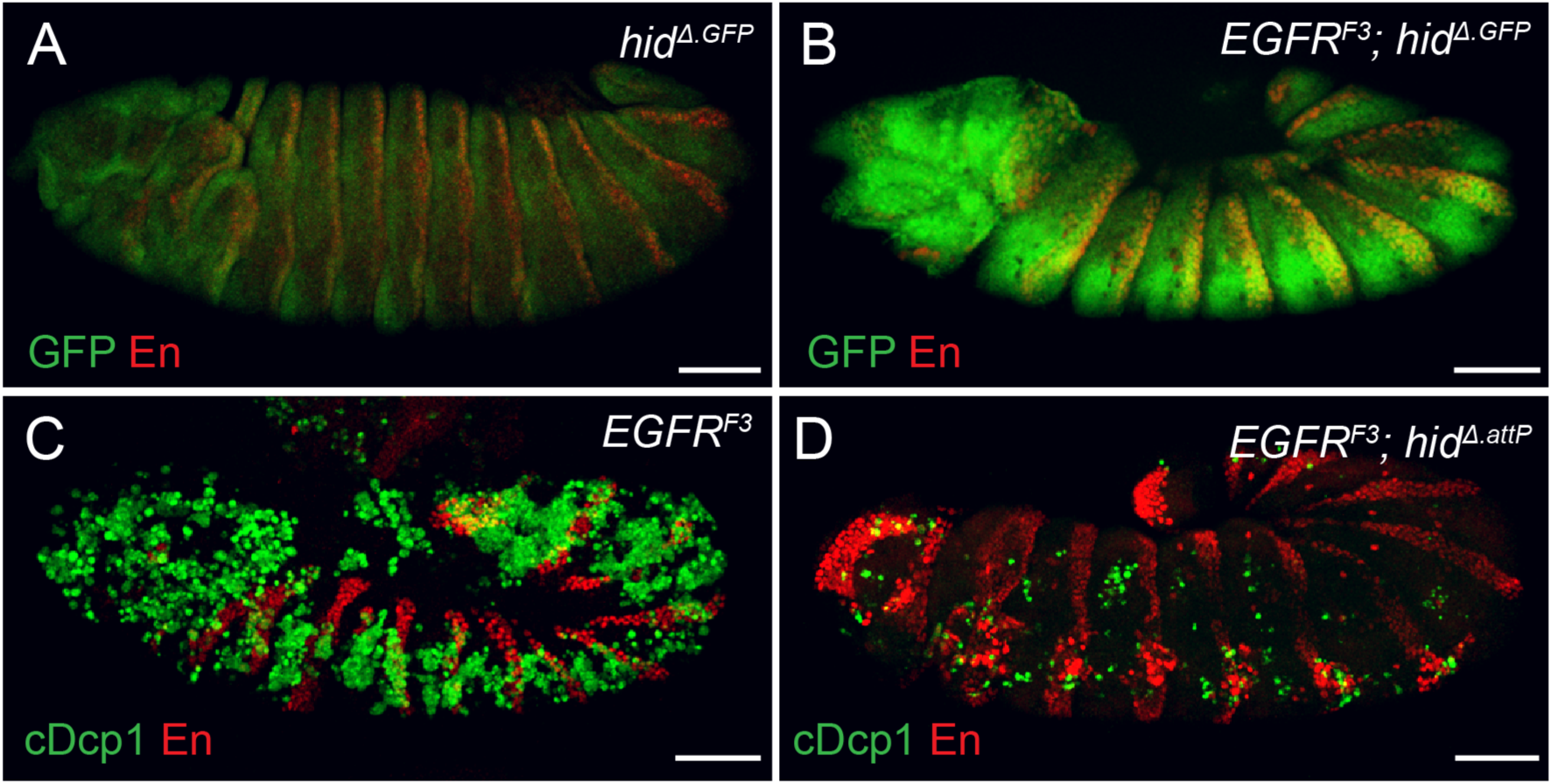
EGFR signalling maintains epidermal cell survival through repression of *hid*. (A, B) Transcription of *hid*, as assayed with the *hid*^*Δ.GFP*^ reporter, in control (A) and *EGFR*^*F3*^ mutant (B) embryos. *hid* is upregulated in most epidermal cells upon loss of EGFR signalling. (C, D) Cleaved Dcp1 immunoreactivity is strongly upregulated throughout the epidermis in *EGFR*^*F3*^ single mutants (C) and this upregulation is lost in *EGFR*^*F3*^; *hid*^*Δ.attP*^ double homozygotes. Scale bars 50µm

**Fig. S4.**
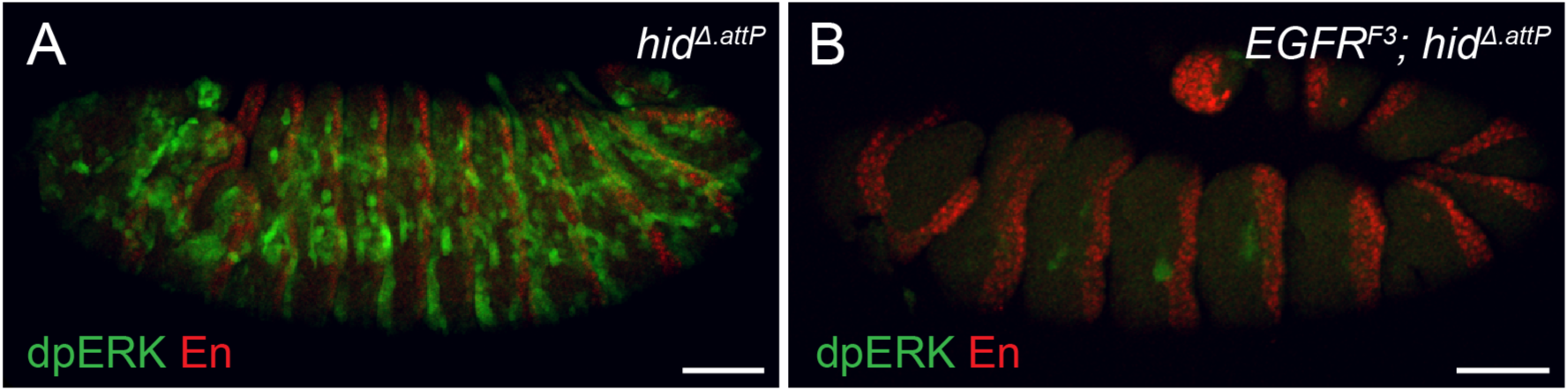
dpERK immunoreactivity is markedly reduced upon loss of EGFR signalling. (A, B) dpERK immunoreactivity in control (*EGFR*^+/+^) *hid*^*Δ.attP*^ (A) and *EGFR*^*F3*^; *hid*^*Δ.attP*^ double homozygotes. Extensive dpERK immunoreactivity is detected in wild type control embryos (A) and this signal is largely lost in *EGFR*^*F3*^; *hid*^*Δ.attP*^ double mutants (B). We take this as evidence that EGFR signalling is the major source of ERK phosphorylation in the embryonic epidermis. Scale bars 50µm.

**Fig. S5.**
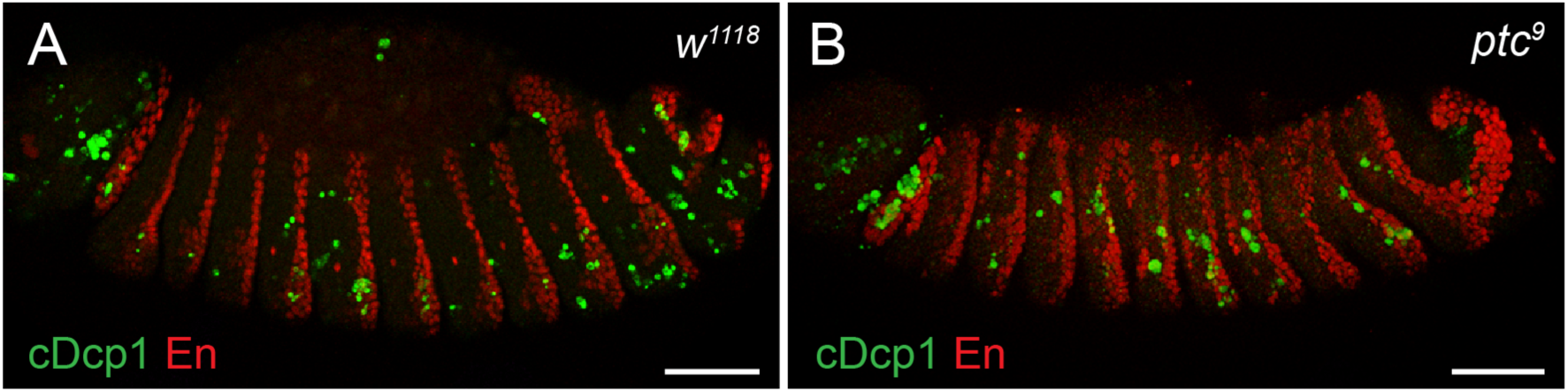
Absence of ectopic apoptosis in *patched* mutants. (A, B) Cleaved Dcp1 immunoreactivity in control *w*^*1118*^ (A) and *ptc*^*9*^ mutant embryos (B). No increase in Dcp1 cleavage was detected in *the ptc* mutants, despite disruption to the segmental pattern. Scale bars 50µm.

**Fig. S6.**
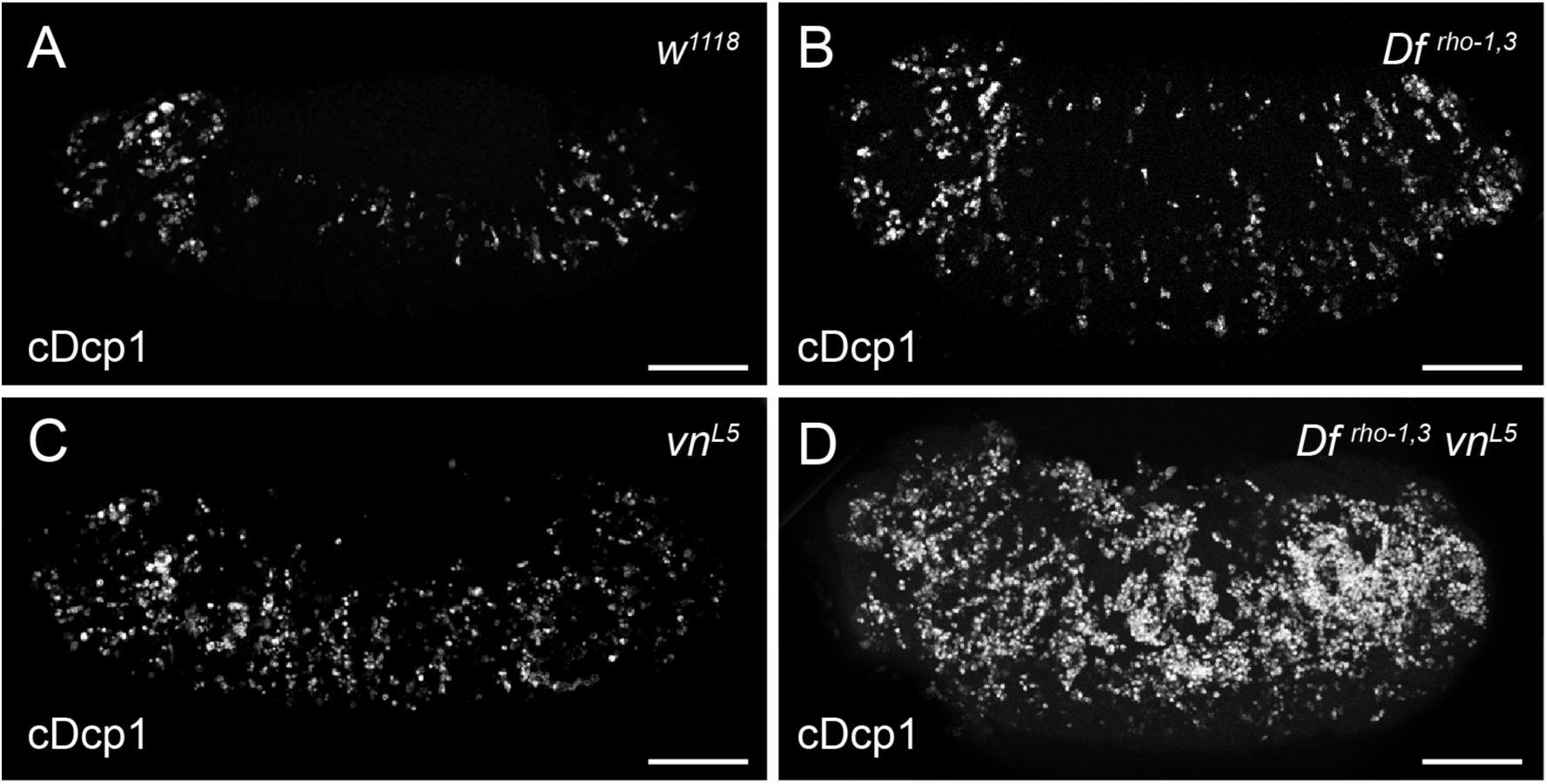
Vein and Rhomboid act redundantly to maintain epidermal cell survival. (A-D) Cleaved Dcp1 in control *w*^*1118*^ (A), *Df* ^*rho*-1,3^ (B), *vn*^*L6*^ (C) and *Df* ^*rho*-1,3^ *vn*^*L6*^ double homozygotes (D). A mild increase in Dcp1 immunoreactivity is seen in *vn* and *rho*-1,3 single mutants (compared to w1118 controls). This signal is strongly enhanced in the double mutants. Scale bars 50µm.

**Table S1.** A genetic survey to identify regulators of apoptosis in patterning mutants List of pathways targeted and transgenes used to assess the rescue of apoptosis in *ftz* mutants. In each instance, the transgene of interest was expressed globally using the *actin5C*-Gal4 driver and a positive score was attributed if a reduction in the levels of cleaved Dcp1 occurred.

| Pathway | Transgene | Apoptotic reduction |
| --- | --- | --- |
| JNK | UAS-*puckered* | - |
| *hedgehog* | UAS-*hh* | - |
| *wingless* | UAS-*wg* | - |
| *dpp* | UAS-*tkv<sup>act</sup>* | - |
| *hippo* | UAS-*yki.GFP* | - |
| p53 | UAS-p53<sup>DN</sup> | - |
| JAK/STAT | UAS-*upd* | - |
| EGFR | UAS-*EGFR* | + |

**Mov. S1.** *hid* is upregulated at mid-embryogenesis in *ftz* mutants Activity of the *hid*^*Δ.GFP*^ reporter in a *ftz*^*Δ.attP*^ mutant embryo. Negligible fluorescence is detected during the early stages of embryogenesis but around embryonic stage 11 (approximately 7 hours after egg laying) bands of fluorescence appears. Hours after egg lay are displayed in the lower right corner.

